# Plasma membrane folding enables constant surface area-to-volume ratio in growing mammalian cells

**DOI:** 10.1101/2024.07.02.601447

**Authors:** Weida Wu, Alice R. Lam, Kayla Suarez, Grace N. Smith, Sarah M. Duquette, Jiaquan Yu, David Mankus, Margaret Bisher, Abigail Lytton-Jean, Scott R. Manalis, Teemu P. Miettinen

## Abstract

All cells are subject to geometric constraints, including the surface area-to-volume (SA/V) ratio, which can limit nutrient uptake, maximum cell size, and cell shape changes. Like the SA/V ratio of a sphere, it is generally assumed that the SA/V ratio of cells decreases as cell size increases. However, the structural complexity of the plasma membrane makes studies of the surface area challenging in cells that lack a cell wall. Here, we investigate near-spherical mammalian cells using single-cell measurements of cell mass and plasma membrane proteins and lipids, which allows us to examine the cell size scaling of cell surface components as a proxy for the SA/V ratio. Surprisingly, in various proliferating cell lines, cell surface components scale proportionally with cell size, indicating a nearly constant SA/V ratio as cells grow larger. This behavior is largely independent of the cell cycle stage and is also observed in quiescent cells, including primary human monocytes. Moreover, the constant SA/V ratio persists when cell size increases excessively during polyploidization. This is enabled by increased plasma membrane folding in larger cells, as verified by electron microscopy. We also observe that specific cell surface proteins and cholesterol can deviate from the proportional size scaling. Overall, maintaining a constant SA/V ratio ensures sufficient plasma membrane area for critical functions such as cell division, nutrient uptake, growth, and deformation across a wide range of cell sizes.

## INTRODUCTION

Surface area-to-volume (SA/V) ratio sets a theoretical maximum for various cell functions, including cell growth, nutrient uptake and shape changes ^1–9^. SA/V ratio is predominantly studied in unicellular organisms, where SA/V ratio can change with cell size and environmental conditions ^10–14^. In animals, cells that require high nutrient uptake or high capacity to deform typically display high SA/V ratio. For example, microvilli on the intestinal enterocytes increase the apical plasma membrane area significantly, which is considered critical for nutrient uptake ^15^. Similarly, T-lymphocytes display high plasma membrane area due to microvilli and membrane folds, which enables cell deformations necessary for tissue intravasation and migration ^16,17^. In the context of cell growth, the SA/V ratio is generally assumed to decrease as cell size increases. Because the ability of cells to take up nutrients can depend on their surface area, the decreasing SA/V ratio could impose an upper limit for cell size and cause size-dependency in cell growth ^7,8,18^. The decreasing SA/V ratio could also enable cell size sensing ^13,19–21^. Yet, little is known about the regulation of SA/V ratio during the growth and division of animal cells.

Changes in the SA/V ratio during cell growth are described by the cell size scaling of cell surface area. This is quantified as the scaling factor (exponent) of the power law:

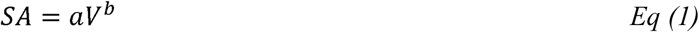

where *a* is a constant and b is the scaling factor. *b* = 1 depicts isometric scaling, where surface area and cell volume grow at the same rate (i.e. SA/V ratio is constant). If *b* ≠ 1, the scaling is called allometric. The specific case of *b*= depicts ⅔-geometric scaling, which is seen in perfect spheres, where the surface area grows at a slower rate than cell volume (i.e. SA/V ratio decreases as cells grow larger). ∼⅔-geometric size scaling is observed in bacteria ^10^ and is expected to apply to animal cells, especially if the cells are nearly spherical in shape. However, this model of size scaling cannot entirely explain cell behavior during proliferation. Proliferation under a steady state requires that cells, on average, double their volume and surface area within each cell cycle, and this requirement is not met by surface area that follows ⅔-geometric size scaling. Here we consider two competing size scaling models for SA/V ratio in proliferating cells that cells may utilize to solve this problem (Fig. 1A).

**Figure 1.**
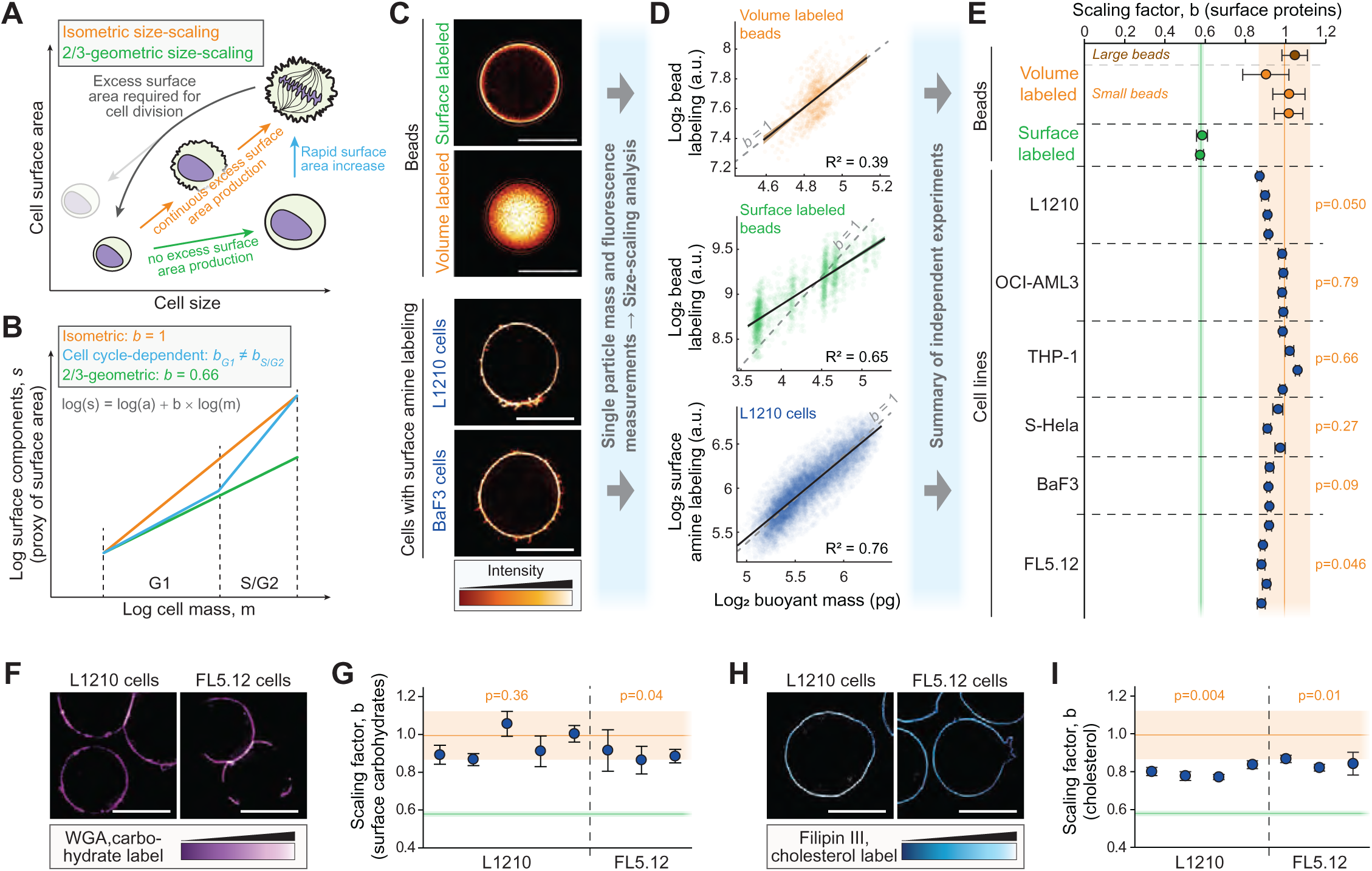
Single-cell mass and surface protein measurements reveal isometric size scaling of plasma membrane proteins in mammalian cells. **(A)** Models for cell size scaling of cell surface area in proliferating cells. Isometric size scaling (orange) would result in cells accumulating surface area at the same rate as volume, resulting in a buildup of plasma membrane reservoirs as cells grow larger. These membrane reservoirs are required for cytokinesis, where the apparent surface area increases. Alternatively, cells could follow ⅔-geometric size scaling of surface area (green), where the membrane reservoirs do not increase during growth, but additional membrane is added at the end of the cell cycle to support cytokinesis (blue). **(B)** The models in panel (A) can be distinguished by examining the power law scaling relationship between cell surface components and cell mass. **(C)** Representative single z-layer images of volume and surface labelled polystyrene beads, and of surface protein labeled L1210 and BaF3 cells. Scale bars denote 10 μm. **(D)** Scatter plots displaying the scaling between bead labeling and bead mass (*top & middle*), and between L1210 cell surface protein labeling and cell mass (*bottom*). Each opaque point represents a single bead/cell, black lines and shaded areas represent power law fits and their 95% confidence intervals, and dashed grey lines indicate isometric scaling (n>1000 beads/cells per experiment). **(E)** Quantifications of the scaling factor, b. Two different sized volume-labeled beads were used as positive controls for isometric scaling. **(F)** Representative single z-layer images of surface carbohydrate labeled L1210 and FL5.12 cells. Scale bars denote 10 μm. **(G)** Quantifications of the scaling factor, b, for cell surface carbohydrate labeling. **(H, I)** Same as (F) and (G), but for Filipin III labeling of cholesterol. In panels (E, G, and I), each dot represents a separate experiment (n>450 beads/cells per experiment), error bars represent the 95% confidence interval, p-values depict comparison to volume labeled beads, and orange and green areas indicate 95% confidence intervals of volume and surface-labeled bead data, respectively.

SA/V ratio could be cell-cycle dependent, so that cells display ⅔-geometric scaling of surface area during most of the cell cycle. In this case, the excess surface area needed for cell division would need to be produced at the end of each cell cycle, resulting in a scaling factor of >1 in during this period (Fig. 1B). This model is supported by the findings that the translation of lipid synthesis enzymes is upregulated at the end of the cell cycle and several lipid species accumulate from S-stage to M-stage ^22–27^. Alternatively, cells could produce volume and plasma membrane components at the same rate, resulting in a constant SA/V ratio, i.e. isometric size scaling of the plasma membrane components (scaling factor of 1, Fig. 1B). This model is supported by the observation that most cell organelles, transcripts and proteins scale isometrically with cell size ^20,28–42^.

Quantifying the area of the plasma membrane by imaging is difficult due to the various membrane folds and nanometer-scale structures ^43^. This has made studies of the SA/V ratio challenging in animal cells, where surface area is not defined by a rigid cell wall. Here, we overcome this challenge by quantifying cell surface localized plasma membrane components as a proxy for surface area, which we then compare to the size of the cells. This allows us to examine the size scaling and cell-cycle regulation of the SA/V ratio, as well as individual cell surface proteins, in mammalian cells. We reveal that cells exhibit approximately constant SA/V ratio as they grow larger. This causes cell size dependency in plasma membrane morphology and enables cell divisions and growth at all sizes. We propose that maintaining a constant SA/V ratio simplifies the regulation of surface area, resulting in a more robust cellular design.

## RESULTS

### Isometric size scaling of plasma membrane associated components in published data

We started by examining the production of plasma membrane associated components using a previously published dataset that links single-cell mass measurements with single-cell RNA-seq (Fig. S1A) ^44^. To obtain statistically rigorous size scaling behaviors, we grouped genes together based on their GO-term association. On average, all cell transcripts displayed scaling factors of 0.91±0.001 and 0.85±0.001 (mean±SE) in L1210 (mouse lymphocytic leukemia) and FL5.12 (mouse pro–B lymphocytes) cells, respectively (Fig. S1B). In contrast, cell division associated transcripts super-scaled with cell size, validating that non-isometric scaling behaviors can be identified. However, transcripts associated with the external side of plasma membrane were not statistically different from all cell transcripts, as the plasma membrane associated transcripts displayed scaling factors of 0.87±0.07 and 0.88±0.04 (mean±SE) in L1210 and FL5.12 cells, respectively. Thus, the cell size scaling of plasma membrane associated mRNAs is near-isometric. This is consistent with the size scaling of most transcripts across various model systems ^20,33,34,36–40^, and similar to the size scaling of plasma membrane associated proteins in human cells ^30^. However, even if plasma membrane proteins are synthesized isometrically with cell size, they may not be inserted into the plasma membrane, as most plasma membrane components also localize to intracellular membranes ^45^. We therefore focused on examining the abundance of components specifically on the cell surface.

### Proliferating mammalian cells display near-isometric size scaling of cell surface components

We established an approach for quantifying the abundance of cell surface proteins as a proxy for cell surface area, as ∼50% of the plasma membrane consists of proteins ^46^. Our approach couples the suspended microchannel resonator (SMR), a cantilever-based single-cell buoyant mass sensor ^47,48^, with a photomultiplier tube (PMT) based fluorescence detection setup ^49^. This enables a measurement throughput of 30,000 single cells/hour (Figs. S2A-B) ^50,51^. We validated our approach’s sensitivity to distinguish different modes of size scaling by measuring spherical polystyrene beads that were labeled either throughout the volume of the beads or specifically on the surface of the beads (Fig. 1C). Volume-labelled beads displayed a scaling factor of 0.99±0.06 (mean±SD) and surface-labelled beads displayed a scaling factor of 0.58±0.01 (mean±SD) (Figs. 1D & E). Notably, the beads had little size variability, indicating that our approach can separate distinct size scaling behaviors even over small size ranges.

Our scaling analyses are carried out using cell buoyant mass as an indicator of size, but many scaling relationships, including SA/V scaling, use cell volume as the size indicator. However, buoyant mass is an accurate proxy for cell volume (Fig. S2C) ^50^ and dry mass ^52^. Therefore, our size scaling analyzes are reflective of both volume scaling and dry mass scaling.

To label cell surface proteins across the external side of the plasma membrane in live cells, we utilized a cell impermeable, amine reactive dye that is coupled to a fluorophore ^53^. Cell labeling was carried out on ice for 10 min to prevent plasma membrane internalization, and the surface-specificity of the labeling was validated with microscopy (Figs. 1C, S2D). Dead cells were excluded from final data analysis (Fig. S2E). As model systems for our work, we focused on suspension grown mammalian cell lines: L1210, BaF3 (mouse pro-B lymphocyte), S-HeLa (suspension grown human adenocarcinoma), OCI-AML3 (human myeloid leukemia), THP-1 (human monocytic leukemia) and FL5.12. These cells maintain near-spherical shape during growth (Figs. 1C, S2D), which makes these cells likely candidates to exhibit ⅔-geometric scaling.

We found that the size scaling of surface protein content was distinct from the surface-labeled beads and similar to the volume-labeled beads in all cell lines (Figs. 1D & E). For example, L1210 and THP-1 cells displayed scaling factors of 0.90±0.02 and 1.01±0.04 (mean±SD), respectively. The scaling factors were statistically different from the volume-labeled beads only for L1210 and FL5.12 cells (Fig. 1E). Pearson correlation values for the fitted scaling factors were high, R^2^ = 0.66±0.08 (mean±SD across all samples), and we verified that our results were not sensitive to outliers or the gating strategy (Figs. S2F-H). We also utilized an alternative cell surface protein labeling chemistry, where protein thiol groups are labeled using fluorescent coupled Maleimide ^53^. This approach also yielded near-isometric scaling factors (Figs. S2I-K). Overall, these results are consistent with the size scaling of mRNAs associated with the external side of plasma membrane (Fig. S1B).

To support our analysis, we also examined the size scaling of other membrane components. We labeled cell surface carbohydrates and cholesterol in L1210 and FL5.12 cells using fluorescent wheat germ agglutinin (WGA) and Filipin III, respectively. We validated the surface specificity of the labeling approaches with microscopy (Figs. 1F, 1H) and analyzed size scaling using the SMR. The cell surface carbohydrates displayed near-isometric size scaling with scaling factors of 0.95±0.08 (mean±SD) in L1210 cells and 0.89±0.03 (mean±SD) in FL5.12 cells (Fig. 1G). In contrast, the cholesterol labeling displayed lower scaling factors of 0.80±0.03 (mean±SD) in L1210 cells and 0.84±0.02 (mean±SD) in FL5.12 cells (Fig. 1I). Thus, cholesterol displayed allometric (sub-isometric) size scaling that was distinct from both surface and volume-labeled beads, as well as from cell surface proteins. However, we note that this cholesterol size scaling was analyzed in fixed cells, where original cell volume may not correlate perfectly with the measured buoyant mass.

Overall, when analyzing across all cells in a population, proliferating mammalian cells exhibit a scaling of surface components that is closer to isometric than ⅔ geometric scaling despite a seemingly spherical cell shape. However, individual surface components can still deviate from isometric scaling, as seen with cholesterol. From here on, we focus on cell surface protein labeling as a proxy for total surface components.

### Cell cycle-specific effects in cell surface protein content

One way for cells to achieve isometric scaling at the population level is if the surface protein content varies with the cell cycle (Fig. 1B). To test this, we first separated G1 and S/G2 stages of the cell cycle using the fluorescent ubiquitination-based cell cycle indicator, geminin-GFP (Figs. 2A, S3A). We observed cell cycle-dependency in the size scaling of surface protein content in L1210, OCI-AML3 and FL5.12 cells, where G1 cells displayed higher scaling factors than S/G2 cells (Fig. 2B). However, all these scaling factors remained above 0.8. To explore this further, and to reveal more gradual changes in the scaling factors, we analyzed the scaling factors in a continuous moving window across different sized cells. This revealed that the scaling factor changes non-monotonically in all cell lines (Figs. 2C, S3B). The scaling factor increased when moving from small to large G1 cells, decreased during the S-phase, and increased again in the largest cells (G2/M). This suggests that cells alter their production of surface proteins and internal components proportionally, with more internal components being produced during S-phase. By examining cells across different ploidy (Fig. S3C), we validated that these surface protein production dynamics could be attributed to cell cycle rather than nonlinear cell size dependent effects.

**Figure 2.**
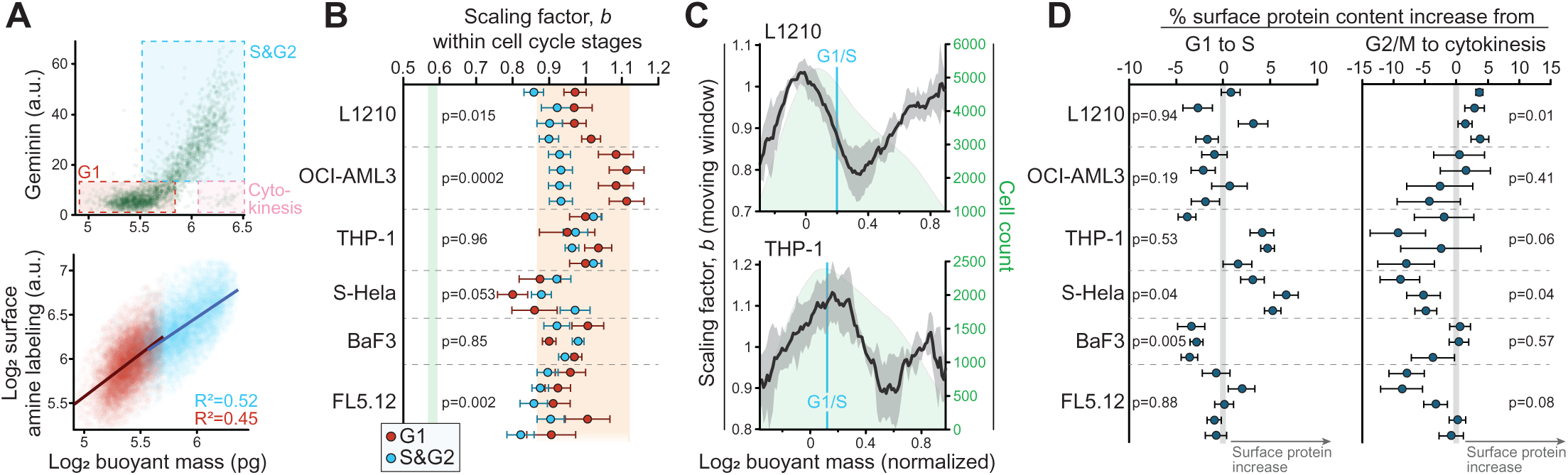
Cell cycle-dependent effects on cell surface protein content. **(A)** *Top*, the gating strategy used to separate cell cycle stages according to cell mass and geminin-GFP. *Bottom*, a scatter plot displaying the scaling between L1210 cell surface protein labeling and cell mass in G1 (red) and S&G2 (blue) stages. Each opaque point represents a single cell, lines represent power law fits to each cell cycle stage. **(B)** Quantifications of the scaling factor, b, in G1 (red) and S/G2 (blue) stages. Each dot represents a separate experiment, error bars represent the 95% confidence intervals (n>500 cells per experiment). Orange and green areas indicate 95% confidence intervals of volume and surface-labeled bead data, respectively. p-values were obtained using Student’s paired t-test between G1 and S/G2 scaling factors. **(C)** Moving window analysis of the scaling factor, b, as a function of cell mass in indicated cell lines. The scaling factor varies within the cell cycle, but does not decrease to ⅔. Dark line and shaded areas represent the moving mean±SD of the scaling factor (N=4 independent experiments), green area depicts the underlying cell size distribution, blue vertical line indicates typical G1/S transition size. **(D)** Cell surface protein content changes between identically sized G1 and S stage cells (left) or G2/M and cytokinetic cells (right). Each dot represents a separate experiment, error bars represent the compound SEM. p-values were obtained using Student’s t-test and reflect comparisons to 0.

Cell cycle transitions could impose rapid changes on cell surface contents. For example, cell divisions impose additional membrane requirements on the cells, which are estimated to be ∼30% in a typical lymphocyte ^53^, and cells could meet this membrane requirement by a rapid expansion of membrane components during cytokinesis. To directly investigate this, we compared the cell surface protein content between identically sized cytokinetic (geminin-GFP negative) and G2/M (geminin-GFP positive) cells ^53^. We did not observe differences between G2/M and cytokinetic cells that would be systematic across all cell lines, and most cells did not increase their surface protein content when entering cytokinesis (Fig. 2D). Thus, the increased membrane requirement imposed by cytokinesis is unlikely to be met by membrane addition taking place specifically during cell division ^53^. We also investigated the G1/S transition using a similar approach, but we did not observe systematic changes shared by all cell lines (Fig. 2D).

Together, these results show that cells i) do not adopt ⅔-geometric size scaling of their surface components even in specific cell cycle stages and ii) do not radically alter their surface protein content at cell cycle transitions. However, cell cycle progression is still associated with changes in the relative production of cell surface proteins and internal components. For L1210 cells, changes in the relative production of cell surface proteins and internal components mirror the cell growth rates (Figs. S3C&D) ^54^. We also note that S-HeLa cells displayed different cell cycle-dependencies than other cell lines used in our study (Figs. 2B, 2D, S3B). This may reflect cell differentiation, as our other model systems originate from hematopoietic lineage.

### Individual cell surface proteins display heterogenous size scaling and cell cycle-dependency

The cell size scaling of individual proteins could differ from the collective size scaling of all cell surface proteins, as shown for many intracellular proteins ^28,30^ and the plasma membrane localized cell polarity regulating PAR proteins ^55^. We selected 4 highly expressed proteins with a cluster of differentiation identifier (CD number) from our scRNA-seq data and used immunolabelling of live cell surfaces to analyze the size scaling of the individual proteins. Surface-specificity of the labeling was validated using microscopy (Fig. 3A, top). The proteins examined included amino acid transporters (Solute Carrier Family 3 Member 2 (SLC3A2), a.k.a. CD98), membrane receptors and signaling proteins (B-Lymphocyte Surface Antigen B4 (CD19); and Protein Tyrosine Phosphatase Receptor Type C (PTPRC), a.k.a. CD45), and cell adhesion molecules (Intercellular Adhesion Molecule 1 (ICAM1), a.k.a. CD54). Curiously, each protein displayed a distinct size scaling behavior (Fig. 3A). For example, CD19 displayed near-isometric size scaling (scaling factor 0.92±0.03, mean±SD), which was identical to the overall surface protein labeling. In contrast, SLC3A2 displayed lower scaling factors (scaling factor 0.73±0.03, mean±SD) and a cell cycle-dependency where S&G2 cells displayed ⅔-geometric size scaling (scaling factor 0.58±0.04, mean±SD). Curiously, this SLC3A2 size scaling, which reflects only the surface localized protein scaling, is distinct from that previously reported for SLC3A2 in experiments that measured total proteome size scaling ^30^. ICAM1 displayed a cell cycle-dependency where the scaling factors were high in G1 (scaling factor 0.98±0.15, mean±SD) but significantly lower in S/G2 (scaling factor 0.76±0.06, mean±SD). These results suggest the existence of cell size-dependent amino acid uptake and cell adhesion specifically in S/G2 cells. More broadly, these results highlight that individual cell surface proteins are regulated independently of each other, resulting in heterogeneous size scaling and cell cycle-dependency.

**Figure 3.**
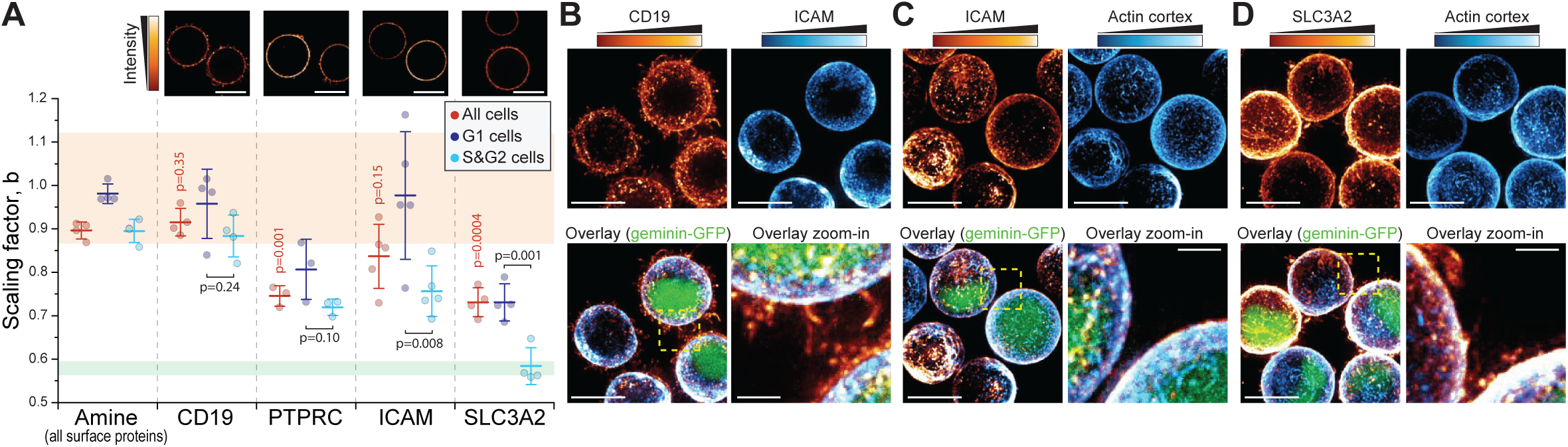
Individual cell surface proteins display heterogeneity in their cell size scaling. **(A)** *Top*, representative single z-layer images of L1210 cell surface immunolabeling (N=2-3 independent experiments, n>40 cells). *Bottom*, quantifications of the scaling factor, b, for indicated cell surface proteins. Overall surface protein (amine) labeling is shown for reference. Data is shown for whole population (red), and separated G1 (dark violet) and S&G2 (light blue) stages. Each dot represents an independent experiment (n>500 cells), bars and whiskers represent mean±SD. Orange and green areas indicate 95% confidence intervals of volume or surface labeled bead data, respectively. p-values in red indicate Welch’s t-test between individual and overall protein labeling. p-values in black indicate Student’s paired t-test between G1 and S&G2 data for individual proteins. **(B-D)** Representative maximum intensity projections of immunolabelled live L1210 cells. The immunolabeling was compared to the LifeAct F-actin sensor (C-D), except in (B) where the comparison was to ICAM immunolabeling. Scale bars denote 10 μm, except in zoom-ins, where scale bars denote 2 μm. N=2-3 independent experiments and n>55 cells.

Next, we considered the mechanistic basis for the heterogeneity of scaling factors between individual plasma membrane proteins. Many plasma membrane proteins bind to the actin cortex, but folding of the plasma membrane spatially separates parts of the plasma membrane from the actin cortex. Consequently, the space available for plasma membrane proteins to bind the actin cortex could display ⅔-geometric scaling. We therefore examined if the plasma membrane proteins with ⅔-geometric size scaling are exclusively colocalized with the actin cortex, whereas the membrane proteins with isometric size scaling are found also on membrane folds. We immunolabeled CD19, ICAM1 and SLC3A2 in L1210 cells expressing the LifeAct F-actin reporter. CD19, which displayed isometric size scaling, was clearly present on membrane folds and ruffles (Fig. 3B). In contrast, ICAM1, which displayed cell cycle-dependent size scaling, was excluded from the membrane folds and ruffles (Fig. 3B-C), as expected based on the protein’s known association with the actin cortex ^56^. However, we did not observe cell cycle-dependency in ICAM1’s localization, which argues against our hypothesis. Finally, SLC3A2, which displayed ⅔-geometric size scaling in S&G2 cells, was present on membrane folds and ruffles even in S&G2 cells (Fig. 3D). Thus, connection to the actin cortex does not explain the size scaling of individual plasma membrane proteins, suggesting that actin-binding membrane proteins are not limited by the space available on the actin cortex.

### ⅔-geometric size scaling of surface proteins does not arise following cell cycle exit

The isometric size scaling of plasma membrane protein content may be necessary for satisfying requirements imposed by cell divisions, as cells typically need to double their plasma membrane content to divide (Fig. 1A). If so, then non-proliferating cells could display ⅔-geometric size scaling of surface components (Fig. 4A). To examine the size scaling of plasma membrane proteins in the absence of cell divisions and cell cycle progression, we first studied FL5.12 cells that have entered quiescence due to IL-3 starvation ^57^, and THP-1 cells that have entered senescence due to treatment with CDK4/6 inhibitor Palbociclib ^30^. Following a treatment to stimulate cell cycle exit, both cell lines displayed G1 arrest, as evaluated based on DNA content and Geminin-GFP protein levels (Figs. S4A&B). The models also exhibited cell size changes. Quiescence resulted in cell volume decrease, and senescence resulted in cell volume increase, when compared to proliferating control populations (Figs. S4C&D). Cell density decreased in senescent THP-1 cells, as seen in other senescent models ^58^, but cell density increased in quiescent FL5.12 cells (Figs. S4E&F). Both model systems retained near-spherical morphology following the cell cycle exit (Fig. 4B). In FL5.12 cells, cell cycle exit resulted in an increased scaling factor (1.15±0.17, mean±SD) when compared to proliferating control cells (0.90±0.03, mean±SD) (Figs. 4C, top). In THP-1 cells, cell cycle exit resulted in a scaling factor that was decreased in comparison to proliferating control cells (Figs. 4C, middle). However, the scaling factor in senescent THP-1 cells remained above 0.8 and thus significantly different from the ⅔ geometric size scaling. To support our findings in a more physiologically relevant model system, we studied CD14+ primary human monocytes. These cells are terminally differentiated and thus non-proliferative, yet approximately spherical in their morphology (Fig. 4B, bottom). The size scaling of plasma membrane protein content in the monocytes displayed significant sample-to-sample variation with an average scaling factor of 1.33±0.41 (mean±SD) (Fig. 4C, bottom). This scaling was not statistically different from the volume labeled beads (p=0.20, Welch’s t-test, see Supplementary Discussion). Overall, these results indicate that cell proliferation, and the membrane requirements imposed by cell divisions, do not explain the isometric size scaling of plasma membrane contents. Instead, the isometric size scaling of plasma membrane contents may reflect a more fundamental cellular organization principle.

**Figure 4.**
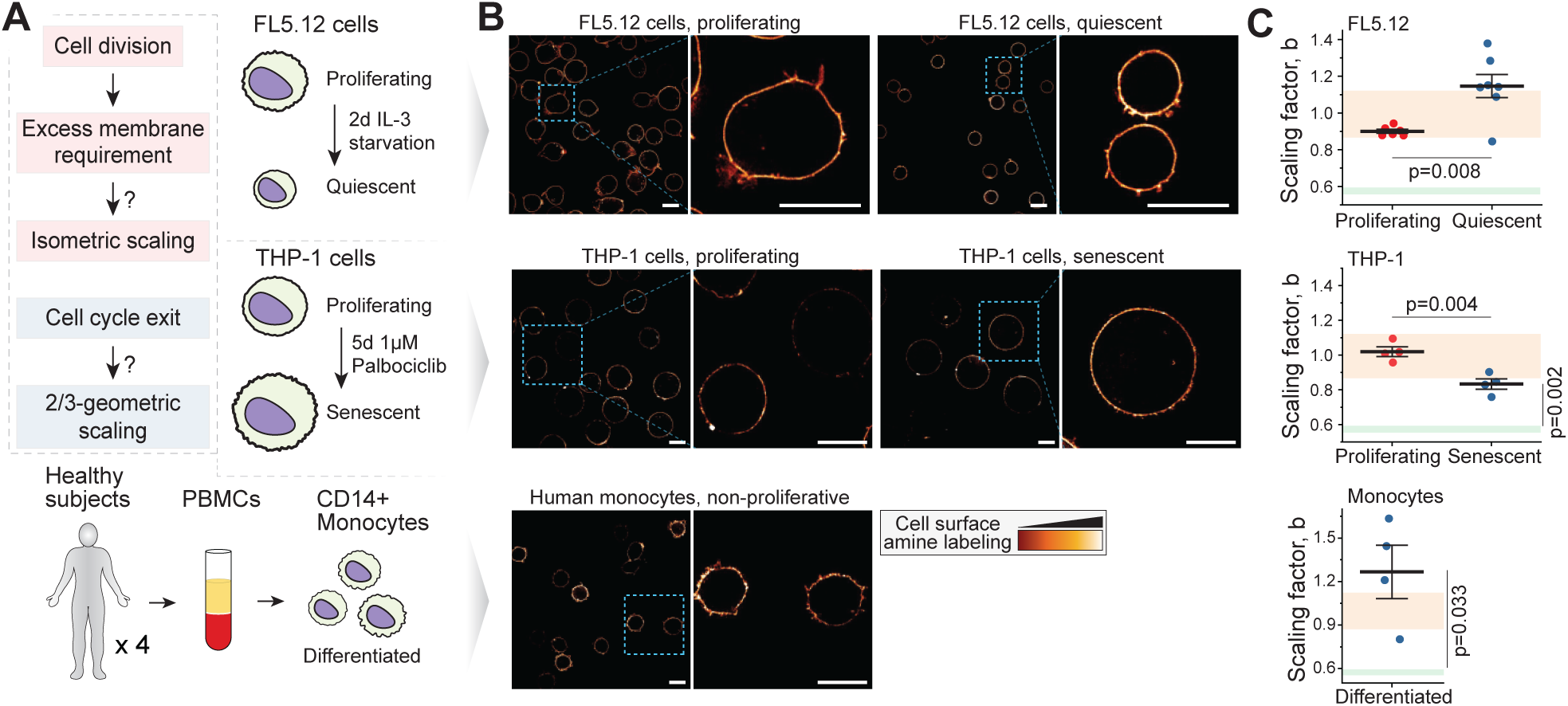
Near-isometric size scaling of surface proteins is not limited to proliferating cells. **(A)** Experimental hypothesis (*left*) and model systems (*right & bottom*). In proliferating cells, plasma membrane content approximately doubles every cell cycle to enable cytokinesis. Upon cell cycle exit, such requirement no longer exists, and the size scaling of cell surface components could follow ⅔-geometric scaling. **(B)** Representative single z-layer images of cell surface amine labeling in proliferating and non-proliferating FL5.12 cells (*top*) and THP-1 cells (*middle*), and in non-proliferating primary human monocytes (*bottom*). Scale bars denote 10 μm. N=2 independent experiments, n>30 cells. **(C)** Quantifications of the scaling factor, b. The high noise in monocyte scaling factors can arise from monocyte isolation or biological patient-to-patient variability. Each dot represents a separate experiment (N=6-7 for FL5.12, N=4 for THP-1 and monocytes), bar and whiskers represent mean±SEM. p-values were obtained using Welch’s t-test when comparing proliferation states to each other and using Student’s one sample t-test when comparing non-proliferating cells to ⅔ geometric size scaling.

### Excessive cell size increases do not downregulate cell surface expansion

We next considered the possibility that the isometric size scaling of plasma membrane protein content is only maintained when cells are within a normal size range. Following excessive cell size increases, feedback mechanisms may decrease the rate of plasma membrane expansion to prevent excessive plasma membrane accumulation (Fig. 5A). We therefore compared a model where cells adjust their surface area expansion following excessive cell size increases to a model where size scaling of surface area remains constant at all sizes. We treated L1210 and THP-1 cells with Barasertib, an Aurora B inhibitor which prevents cytokinesis, to generate polyploid cells with large cell size increases ^54^. We then combined cell populations at different stages of polyploidization, labelled cell surface proteins and analyzed the cells for size scaling (Figs. 5B-D). In both L1210 and THP-1 cells, despite ∼10-fold increases in cell size, the surface protein size scaling was independent of cell size and ploidy (Figs. 5E&F). Thus, we did not observe feedback mechanisms that would decrease the rate of plasma membrane expansion as cells grow excessively large.

**Figure 5.**
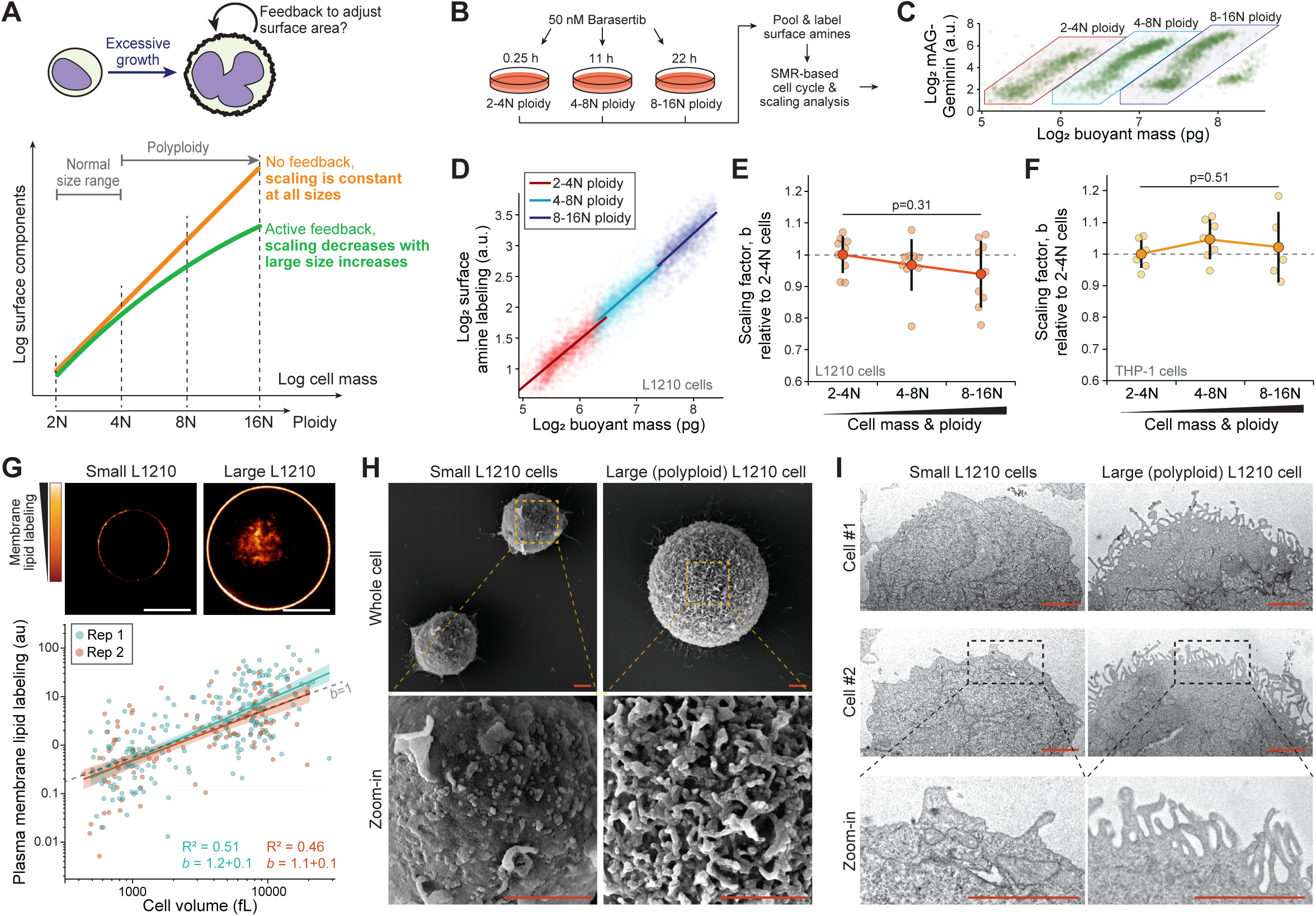
Excessive cell growth is coupled with plasma membrane folding and isometric size scaling of cell surface area. **(A)** Schematic of competing models. If cells aim to maintain a fixed degree of plasma membrane folding, excessive cell size increases should decrease the scaling factor (green). However, if cells produce surface components independently of the degree of plasma membrane folding, the scaling factor should be independent of cell size (orange). **(B)** Experimental setup. **(C)** A scatter plot of geminin-GFP as a function of cell mass in L1210 cells of different ploidy. **(D)** A scatter plot displaying the scaling between cell mass and surface amine labeling in L1210 cells. Each opaque point represents a single cell colored according to its ploidy. Power law fits are shown separately for each ploidy (n>1000 cells per ploidy). **(E)** Quantifications of the scaling factor, b, across L1210 cells of different ploidy. Each small, opaque point indicates an independent experiment (N=9). Large, dark points and error bars represent mean±SD. p-value obtained using ANOVA. **(F)** Same as panel (E), but data is for THP1 cells (N=7). **(G)** *Top,* representative single z-layer images of membrane lipid labeling. Scale bars denote 10 μm. *Bottom,* a scatter plot displaying the scaling between cell volume and plasma membrane lipid labeling in L1210 cells of various ploidy. Each opaque point represents a single cell, power law fits are shown separately for each experiment (N=2 independent experiments, n=190 and 124 cells). **(H)** Representative SEM images of small and large (polyploid) L1210 cells. Scale bars denote 2 μm. N=3 independent experiments, n=63 cells. **(I)** Representative TEM images of small and large (polyploid) L1210 cells. Scale bars denote 2 μm. N=2 independent experiments, n=65 cells.

Next, we examined the size scaling of plasma membrane lipids to ensure that our results are not limited to the protein content of the membrane. Our SMR-based approach in polyploid L1210 cells revealed cholesterol size scaling factor of 0.8 (Fig. S5A). This is consistent with the cholesterol size scaling in freely proliferating cells (Fig. 1I). To analyze additional lipids on the plasma membrane, we utilized a generic lipid labeling approach that also labels cell’s internal membranes. Because our SMR-based approach does not differentiate cytoplasmic and plasma membrane labeling, we used microscopy to analyze the plasma membrane-specific lipid labeling in freely proliferating (small) and polyploid (large) L1210 cells (Fig. 5G). The wider dynamic range of the microscopy-based approach also allowed us to study more extreme polyploidization which increased cell volumes nearly 100-fold. The cells maintained near-spherical shapes at all sizes. We observed that the L1210 cells scale their plasma membrane lipids nearly isometrically with a scaling factor of 1.14±0.12 (mean±SD) (Fig. 5G). Together with our previous findings, these results indicate that, on average, components of the plasma membrane accumulate at an approximately fixed rate as cells grow larger, and that cells do not downregulate their cell surface expansion despite excessive size increases.

### Plasma membrane folds enable a constant SA/V ratio as cells grow larger

Our results suggest that plasma membrane ultrastructure must accumulate folding and structural complexity as cells grow larger. To visualize and qualitatively verify this, we carried out scanning electron microscopy (SEM) of L1210 and THP-1 cells with different levels of ploidy. This revealed that L1210 cell plasma membrane folding was radically higher in large polyploid cells in comparison to small diploid cells (Figs. 5H, S5B). While the small diploid L1210 cells displayed large areas of relatively smooth membrane, the large polyploid L1210 cells were uniformly covered in highly folded membrane. Similarly, membrane folding and complexity increased with cell size in THP-1 cells (Fig. S5C). However, the membrane folding in large polyploid THP-1 cells was often polarized to a specific region of the cell and took the form of very large folds rather than small uniform protrusions, as seen in L1210 cells. We also examined proliferating and quiescent FL5.12 cells, as well as proliferating and senescent THP-1 cells using SEM. The small (quiescent) FL5.12 cells were largely devoid of any large membrane folds, which were common on the normal-sized (proliferating) FL5.12 cells (Fig. S4G). Similarly, large membrane folds were more abundant on large (senescent) THP-1 cells than on normal-sized (proliferating) THP-1 cells (Fig. S4H). These results suggest that the amount of membrane folding is dependent on cell size rather than the proliferative state. To verify that our results were not specific to the SEM approach, we also examined small and large (polyploid) L1210 cells using transmission electron microscopy (TEM). This revealed that the plasma membrane of the large polyploid cells adopts microvilli-like structures which were not observed in the small L1210 cells (Fig. 5I). Overall, these results support our conclusion that plasma membrane area scales approximately isometrically with cell size, which causes increased membrane folding in larger cells.

## DISCUSSION

Human cell types vary in size by 7 orders of magnitude ^59^ and, even within a single cell type, proliferating cells exhibit at least a two-fold size variation. Regardless of size, cells must maintain sufficient plasma membrane area to deform, uptake nutrients and interact with their environment. However, directly measuring plasma membrane area (in µm²)—and thus the SA/V ratio—is challenging because the results depend on imaging resolution. As measurement precision increases, finer surface details become visible, leading to progressively larger estimates of membrane area, analogous to the *coastline paradox* in two-dimensional systems ^43^. By instead analyzing how plasma membrane components scale with cell size, we circumvent this issue, providing a robust proxy for estimating SA/V scaling. However, our approach has limitations: we cannot measure all plasma membrane components simultaneously, nor can we precisely weigh the contributions of different components, which may introduce variability in the estimated SA/V scaling.

Our findings indicate that the SA/V ratio remains approximately constant during cell growth. This aligns with previous studies showing that cell size scaling of most transcripts, proteins and cell organelles is near-isometric ^20,28–42^. This near-constant SA/V ratio persists despite nearly a 100-fold increase in cell size. Using the same model system, we have previously demonstrated that cells can sustain exponential cell growth even at these enlarged sizes ^54^. Together, these results suggest that the cell surface area does not become a limiting factor for cell growth as mammalian cells grow larger. Notably, our study focused on near-spherical cells where the expected ⅔-power geometric scaling of surface area is most likely to occur. Since we do not observe this scaling behavior, it is unlikely to be a general feature of mammalian cells.

A long-standing question in cell biology is ‘how do cells sense their size?’ If SA/V ratio decreased with increasing cell size, it could serve as a metric for cells to sense their size ^13,19,20^. Our results argue that mammalian cells do not use the SA/V ratio for cell size sensing. However, more specific molecular mechanisms could still operate in an analogous manner ^55^. We have identified that cholesterol and specific cell surface proteins do not follow isometric size scaling making these components potential cell size sensors. Furthermore, proteins that sense plasma membrane curvature, such as I-BAR proteins and tetraspanins ^21,60,61^, could also act as cell size sensors, because the constant SA/V ratio requires that the plasma membrane accumulates more folding as cell size increases.

The SA/V ratio also impacts cells’ ability to deform ^6^. For example, blood and immune cells frequently pass through the microvasculature by squeezing through constrictions, and this is enabled by plasma membrane reservoirs ^16,17,62^. The constant SA/V ratio may enable immune cells to grow larger, as seen during T and B cell activation ^50^, while undergoing efficient intravasation independently of their size. Curiously, increases in membrane folding are also observed on larger nuclei ^63^, suggesting that nuclear deformation could also be size independent. In addition, the size-dependent plasma membrane morphology that we observe may result in cell size-dependent mechanosensing, endocytosis and exocytosis, all of which can be impacted by local membrane geometry and/or tension ^21,61,64–67^. Notably, as cholesterol can influence membrane elasticity ^68,69^, the mechanical properties of the plasma membrane could also be impacted by the allometric (sub-isometric) cell size scaling of cholesterol.

More broadly, maintaining approximately constant SA/V ratio can simplify surface area regulation. Cell division increases the apparent cell surface area. Should the SA/V ratio decrease as cells grow larger, cells would have to sense how much plasma membrane is needed to facilitate cytokinesis. This is dependent on the cell division size, necessitating either cell size or growth-sensing. In contrast, when the SA/V ratio is maintained constant, there is no need for additional plasma membrane during cytokinesis, and cells do not require cell size or growth sensing to maintain an appropriate cell surface area. This makes cytokinesis robust towards variability in cell division size, while also ensuring that plasma membrane area does not become limiting for any other cell functions due to cell size increases. We propose that the constant SA/V ratio may represent a fundamental cellular design principle that enables cells to robustly grow and operate across a range of cell sizes.

## ACKNOWLEDGEMENTS

SRM and TPM received funding from NIH’s National Institute of General Medical Sciences (R01GM150901). SRM also received funding from the MIT Center for Cancer Precision Medicine and Virginia and DK Ludwig Fund for Cancer Research. The work was supported by the Koch Institute Support (core) Grant P30-CA14051 from the National Cancer Institute, and we thank the Koch Institute’s Robert A. Swanson (1969) Biotechnology Center for technical support, specifically the Peterson (1957) Nanotechnology Materials Core Facility (RRID:SCR_018674) and the Microscopy Core Facility. We also thank Prof. J. Chen and Dr. N. Ivica for providing the wt OCI-AML3 and THP-1 cells, Dr. L. Debaize fron Dana-Farber Cancer Institute for assisting with primary patient samples, and Michael Collis from Spherotech for providing volume-labeled beads.

## AUTHOR CONTRIBUTIONS

W.W., K.S. and T.P.M. carried out SMR and fluorescence microscopy experiments. W.W. analyzed the SMR data. A.R.L. and G.N.S. carried out image analysis. S.M.D. and J.Y. handled primary patient samples. D.M., M.B., A.L-J. and T.P.M carried out electron microscopy. S.R.M. and T.P.M. supervised the study. T.P.M. wrote the manuscript with assistance from W.W., A.R.L., and S.R.M. The study was conceived by T.P.M.

## DECLARATION OF INTERESTS

S.R.M. is a co-founder of Travera and Affinity Biosensors, which develop technologies relevant to the research presented in this work. The other authors declare no competing interests.

## METHODS

### Cell lines and cell cycle reporters

All cells were cultured in a humidified incubator at 37°C under 21% O_2_ and 5% CO_2_. L1210 (mouse, sex = female, obtained from ATCC, #CCL-219), BaF3 (mouse, sex = unknown, BCR-ABL T315I mutated, gifted by the Weinstock lab at Dana-Farber Cancer Institute), OCI-AML3 (human, sex = male, gifted by the Chen lab at MIT), THP-1 (human, sex = male, gifted by the Chen lab at MIT), FL5.12 (mouse, sex = unknown, gifted by the Vander Heiden lab at MIT) and S-HeLa (human, sex = female, gifted by the Elias lab at Brigham And Women’s Hospital) cells were cultured in RPMI (Invitrogen # 11835030) supplemented with 10% FBS (Sigma-Aldrich), 10mM HEPES, 1mM sodium pyruvate and antibiotic/antimycotic. S-HeLa cell cultures were maintained on ultra-low attachment plates (Sigma-Aldrich, #CLS3471-24EA) to prevent cell adhesion. FL5.12 cells were cultured in the presence of 10 ng/ml IL-3 (R&D Systems). Cell cycle exit in FL5.12 was achieved by washing the cells three times with PBS to remove IL-3 from the cells and by placing the cells in IL-3 free culture media for 48 h. All experiments were carried out when cells are at exponential growth phase at confluency of 200,000-700,000 cells/ml. All cell lines tested negative for mycoplasma. Authenticity of all cell lines was evaluated based on cell morphology.

L1210 cells expressing the FUCCI sensor were generated in a previous study ^70^. Other cell lines were transduced with lentiviral vector carrying geminin-GFP encoding plasmid and puromycin resistance (Cellomics Technology, #PLV-10146-200). Transductions were carried out using spinoculation. Approximately 50,000 cells were mixed with ∼2.5*10^6^ virus particles in 400 µl of culture media containing 8 μg/ml polybrene and the mixture was centrifuged for 1 h at 800 g at RT. After centrifugation, cells were resuspended in normal culture media and grown o/n. The spinoculation procedure was repeated the next day after which selection was started using 5 μg/ml of puromycin. Following a week of selection, the cells were sorted for GFP positive cells using BD Biosciences FACS Aria and this was followed by a clonal selection in media containing the selection marker. Clones were evaluated for biphasic geminin-GFP expression using flow cytometry. The top clones used for experiments were also examined for the loss of the geminin-GFP reporter signal at mitosis using timelapse microscopy with the IncuCyte live cell analysis imaging system (Sartorius).

### Primary cells

Primary human monocytes were isolated from apheresis leukoreduction collars obtained from anonymous healthy platelet donors at the Brigham and Women’s Hospital Specimen Bank under an Institutional Review Board–exempt protocol. The donors’ sexes are unknown. First, human peripheral blood mononuclear cells (PBMCs) were isolated using density gradient centrifugation of the collars (Lymphoprep, StemCell Technologies Inc, #07801). Monocytes were then enriched from the PBMC samples via the EasySep Human Monocyte Enrichment Kit (StemCell Technologies Inc, #19059). Monocytes were seeded at a concentration of 500.000 cells/mL in blood cell growth media (Sigma-Aldrich, #615-250) and cultured in a 24 well ultra-low attachment well plates (Sigma-Aldrich, #CLS3473-24EA) at 37°C for 2 h prior to sample staining and measurements. Each experiment was carried out using monocytes isolated from a different patient.

### Scaling control beads

For surface labelled beads, SPHERO Amino Polystyrene beads ranging from 8.0 to 12.9 µm in diameter (Spherotech, #AP-100-10) were labelled with amine labeling as detailed below for cells. For volume labelled beads, we used readily fluorescent (Nile Red) polystyrene beads ranging from 5.0 to 7.9 µm in diameter (Spherotech, #FP-6056-2), which are labeled “small” in our main figure, and we also tested readily fluorescent (CyBlue) polystyrene beads ranging from 8.1 to 12.0 µm in diameter (Spherotech, #FP-10066-2), which are labeled “large” in our main figure. The Nile Red volume labeled beads were FACS sorted using BD Biosciences FACS Aria, to enrich the population for the smallest and largest beads.

### Cell surface labeling approaches

Amine surface labeling of cells was carried out using LIVE/DEAD Fixable Red Dead Cell Stain kit (Invitrogen, #L23102), as detailed previously ^53^. Briefly, the cells were washed with ice cold PBS, mixed in ice cold PBS with the amine reactive stain at 5x supplier’s recommended concentration, and stained in dark on ice for 10 minutes. Staining was stopped by mixing the cells with cold culture media that contains FBS, followed by a washing step with cold PBS. Cells were mixed in PBS at 500.000 cells/ml and immediately analyzed using SMR or microscopy. Thiol surface labeling was carried out as amine labeling, but the stain used was Alexa Fluor 568 C5 Maleimide (Invitrogen, #A20341) at 50 µM concentration. The Alexa Fluor 568 C5 Maleimide stained cells were washed an extra time with cold culture media to remove stain aggregates.

Cell surface carbohydrates and cholesterol were labeled with wheat germ agglutinin (WGA, Invitrogen, #W11263) and Filipin III (Cayman Chemical, #70440), respectively ^53^. For WGA, live cells were labeled in PBS at +4°C for 20 min, whereas for Filipin III, cells were fixed with paraformaldehyde prior to labeling and labeled in PBS at RT for 1 h. Cells were washed twice with PBS after labeling. Plasma membrane lipid labeling in polyploid cells was carried out using CellMask Deep Red Plasma membrane stain (Invitrogen, #C10046) with 1:1000 dye dilution in culture media. Labeling was done at +4°C for 15 min, after which cells were washed twice with PBS, plated and imaged at RT.

Plasma membrane protein immunolabeling was carried out in culture media on ice. First, non-specific Fc receptor binding was blocked using 1:100 dilution of TruStain FcX anti-mouse CD16/32 antibody (BioLegend, RRID:AB_1574975, Cat#101320). After 5 minutes of Fc blocker treatment, antibodies targeting the proteins of interest were added at 1:200 dilution for 15 min. The antibodies used were PE anti-mouse CD19 Antibody (BioLegend, RRID:AB_313643, Cat#115508), APC anti-mouse CD45 Antibody (BioLegend, RRID:AB_2876537, Cat#157606), APC anti-mouse CD54 Antibody (BioLegend, RRID:AB_10612936, Cat#116120) and PE anti-mouse CD98 (4F2) Antibody (BioLegend, RRID:AB_2190813, Cat#128208). For anti-CD98, a APC variant of the same antibody was also used (BioLegend, RRID:AB_2750544, Cat#128211). After immunolabeling, the cells were washed twice with cold culture media and immediately analyzed using SMR or microscopy.

The purity of primary monocyte isolation was assessed using CD14 immunolabeling. The cells were incubated with CD14 Monoclonal Antibody (61D3) conjugated to Alexa Fluor 488 (Invitrogen, eBioscience, RRID:AB_2744748, Cat#53-0149-42) for 20 min at +37°C, and washed twice with PBS. The labelled cells were analyzed using BD Biosciences flow cytometer Celesta with 488 nm excitation laser and 530/30 nm emission filter.

### SMR mass and fluorescence measurements

Cell mass measurements were conducted with SMR devices ^48,71^ coupled with an epi-fluorescence microscope (Nikon LV-UEPI2) for fluorescence measurements (Fig. S2A) ^49,50^. PMTs (Hamamatsu, #H10722-20) were used to measure the emission intensity in five different bandwidths, 438/24, 515/30, 595/31, 678/70, and 809/81 nm. Geminin-GFP intensity and surface protein labeling intensity were measured in 515/30, and 678/70 respectively. The exact specifications for the illumination source, objective, and emission filters are described in ^50^. Analog output and voltage modules (National Instruments, #NI-9263, #NI-9215) were used to communicate with the PMTs. SMR data were collected at a data rate of ∼20 kHz and fluorescence intensity signals from PMTs were collected at a data rate of ∼50 kHz.

A custom MATLAB code was used to process the raw SMR and PMT data, as reported previously ^50^. Fluorescence intensity signals from labeled cells were identified based on positive thresholding, defined by an increase in fluorescence higher than three times the baseline’s standard deviation. Event identification was designed such that if any of the 5 fluorescence channels had a positive event, the intensity of all other channels would be recorded. After independently processing the SMR and PMT data, a custom algorism was used to pair the cell mass and fluorescence signals according to their timestamps ^50^. Each SMR event was expected to have a matching PMT event, with a short time delay due to cell travelling between the measurement locations (Fig. S2B). The pairing algorism computed the time differences between each SMR and PMT event, followed by a manual selection step for a range of acceptable time differences. Events that did not have a unique one-to-one match between the SMR and PMT signals were excluded. Sample preparation and measurements were not blind-controlled or randomized because a single individual was often responsible for all relevant steps.

### Chemical perturbations

Cell cycle exit in THP-1 cells was achieved by treating the cells with 1 µM CDK4/6 inhibitor Palbociclib (Cayman Chemical, #16273) for 5 days. Palbociclib was kept present during surface amine labeling. Chemically induced polyploidy was achieved by treating L1210 FUCCI cells with 50 nM Barasertib (a.k.a. AZD1152-HQPA; Cayman Chemical, Cat#11602) ^54^. To obtain cells with varying sizes and cell cycle states, while avoiding cells that are too large for the SMR microchannels, different Barasertib treatment durations (typically 0.25 h, 10 h, 20 h) were pooled together immediately prior to cell surface amine labeling. For polyploid THP-1 FUCCI cells, 100 nM Barasertib was used, and treatment times were 0.25 h, 24 h, and 48 h. Barasertib treatment was maintained throughout the amine labeling and SMR experiments. For microscopy experiments, L1210 cell Barasertib treatment lasted 48 h and THP-1 cell Barasertib treatment lasted 72 h, after which treated and untreated cells were pooled together for sample preparation in the presence of Barasertib.

### Cell volume measurements

Single-cell volumes were measured using a coulter counter (Beckman Coulter). In short, cells were immersed in PBS with 1:100 dilution, and 1 ml of the solution was measured on the coulter counter using a 100 µm diameter cuvette. In a typical experiment, we measured >2000 cells and particles below the size of 100 fl were excluded.

Single-cell volumes were also measured jointly with cell buoyant masses using the SMR according to a previously detailed fluid switching approach ^47^. In short, cell’s buoyant mass was first measured in normal media in the SMR, after which the cell was immersed in high density media that comprised of normal culture media with 35% OptiPrep density gradient medium (Sigma-Aldrich, Cat#D1556). The cell was then flow back through the SMR cantilever to obtain a measurement in high density media. These two buoyant mass measurements were used to calculate the volume of the cell ^47^, and to correlate cell buoyant mass in normal culture media with cell volume.

### Cell cycle analyses

Cell cycle perturbations were validated using flow cytometry-based detection of DNA content and geminin-GFP levels. Following treatment, cells were washed with PBS and fixed by mixing with ice cold 70% EtOH and incubating o/n. The cells were then washed with PBS and labelled with FxCycle PI/RNase Staining Solution (Invitrogen, #F10797) for 30 min in dark at RT. The cells were analyzed using BD Biosciences flow cytometer LSR II with 488 nm and 561 nm excitation lasers and 515/20 nm and 610/20 nm emission filters.

### Light microscopy and image analysis

All microscopy samples were plated on poly-L-lysine (Sigma-Aldrich, #P8920) coated glass bottom CELLVIEW dishes (Greiner Bio-One, #627975) and imaged at RT using DeltaVision wide-field deconvolution microscope. Imaging was done using standard FITC, PE, Alexa594 and APC filters, a 100x oil-immersion objective and oil with refractive index of 1.516 (Cargille Laboratories). For cell surface amine labeling, z-layers were collected with 0.2 µm spacing covering typically a 4 µm height at the middle of the cell. For cell surface protein immunolabeling, z-layers were collected with 0.25 µm spacing covering typically a 10 μm in height. For plasma membrane lipid labeling, z-layers were collected with 1 µm spacing covering typically 12 µm height at the middle of the cells. Image deconvolution was carried out using SoftWoRx 7.0.0 software. For cell surface protein and antibody labeling approaches, images were processed using ImageJ (version 1.53q).

For plasma membrane lipid labeling, we developed a MATLAB-based image analysis approach. In short, cells were first automatically detected and the z-layer at the middle of each cell was manually selected. Then, line profiles were drawn starting from the center of the cell and ending outside of the plasma membrane. Line profiles that were impacted by neighboring cells were automatically excluded from the analysis. The line profiles from each cell were averaged, aligned to the plasma membrane and used to quantify the total intensity of plasma membrane signal (area under the curve, AUC, in the line profiles). We then established how the one-dimensional AUC quantifications compare to the total plasma membrane content in the cell. To achieve this, we assumed a spherical cell with a membrane volume that can be defined as:

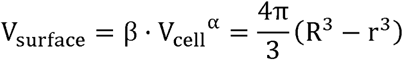

where R depicts cells outer radius, r depicts cells inner radius (to the inner side of the plasma membrane), β is a constant, and α is the scaling factor. The ‘thickness’ of the plasma membrane (T= R - r) corresponds to the AUC of the line profiles. Assuming that α ≈ 1, we can solve for T and obtain

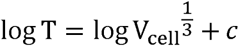

where c depicts a constant. We therefore cubed the quantified line profile AUC values so that isometric size scaling of total plasma membrane corresponds to a scaling factor of 1. For cell size quantifications, we quantified the cell radius from the line profiles and calculated the total cell volume assuming a spherical cell shape.

### Electron microscopy

For SEM, the proliferating and non-proliferating FL5.12 and THP-1 cells were prepared as detailed above and fixed/processed as detailed below. For polyploid experiments, L1210 FUCCI and THP-1 FUCCI cells were treated with Barasertib for 48 h and 72, respectively, at a cell concentration of ∼300.000 cells/ml to induce polyploidy. The polyploid cells and diploid control cells were then mixed in media containing Barasertib. All SEM samples were plated on a round cover slip coated with poly-L-lysine. The cells were given 15 min to adhere, and the media switched to PBS. The cells were then fixed with glutaraldehyde and paraformaldehyde at 2.5% and 2% final concentration, respectively, in PBS for 30 min at RT. After fixation, the cells were washed with PBS and twice with 100 mM sodium cacodylate buffer at 4 °C. Secondary fixation was carried out with 1% osmium tetroxide in the sodium cacodylate buffer for 30 min at 4 °C. The cells were then washed with DI water four times and moved to stepwise dehydration with that was carried out using ethanol (EtOH) at RT. Each step lasted 5 min and the EtOH concentrations were 35%, 45%, 50%, 65%, 70%, 85%, 95% and 100%. Next the cells were treated for 15 min with 50% EtOH and 50% tetramethyl silane (TMS), then 15 min with 20% EtOH and 80% TMS, and finally twice for 5 min with 100% TMS. The TMS was then removed, and the cells were left to dry at RT o/n. Dried SEM samples were sputter coated with gold for 120 s. Imaging was carried out using Zeiss Crossbeam 540 scanning electron microscope at the Peterson (1957) Nanotechnology Materials Core Facility at the Koch Institute for Integrative Cancer Research. Typical imaging was carried out with 5 kV accelerating voltage, 500 pA probe current, a working distance of 8 mm, and a magnification of 4000x. Typical image resolution was 5 nm/pixel.

For TEM, the cells were washed twice with PBS and fixed with glutaraldehyde and paraformaldehyde at 2.5% and 2% final concentrations, respectively, in PBS for 60 min at +4°C. Following fixation, the cells were washed three times with 100 mM sodium cacodylate buffer at 4 °C. Secondary fixation was carried out for 60 min on ice with 1% osmium tetroxide in a solution containing 1.25% potassium ferrocyanide and 100 mM sodium cacodylate. The samples were then washed three times with 100 mM sodium cacodylate and three times with 50 mM sodium maleate at pH 5.2. The samples were stained with 2% uranyl acetate in sodium maleate buffer o/n at RT. The next day, the samples were rinsed twice with distilled H_2_O and dehydrated with ethanol using 10 min incubations at following EtOH concentrations: 30%, 50%, 70%, 95%, 100%, and again 100%. The samples were incubated in propylene oxide for 30 min, twice, and moved to propylene oxide - resin mixture (1:1) for o/n at RT. The following day, the samples were moved to a new propylene oxide - resin mixture (1:2) for 6 hours at RT, and then into full resin o/n at RT. The resin infiltrated samples were moved to molds and polymerized at 60°C for 48 hours. The polymerized resin embedded samples were then sectioned into 60 nm thin slices using a Leica UC7 ultramicrotome and collected on carbon-coated nitrocellulose film copper grids. Imaging was carried out using FEI Tecnai T12 transmission electron microscope at the Peterson (1957) Nanotechnology Materials Core Facility at the Koch Institute for Integrative Cancer Research. Typical imaging was carried out with 120 kV voltage, and a magnification of 4800x, resulting in an image resolution of 3.8 nm/pixel. Images were captured with an AMT XR16 CCD camera. Sample preparation and measurements were not blind-controlled or randomized because a single individual was often responsible for all relevant steps.

### Gene expression data analysis

All gene expression data were obtained from a previous publication ^44^, where SMR-based cell mass and mass accumulation rate measurements were coupled to single-cell collection and Smart-Seq2-based scRNA-seq. The transcriptomic data represents normalized mRNA count measurements (transcripts per million), which was considered to reflect mRNA concentrations in each cell. These mRNA concentrations were converted to a proxy of absolute mRNA levels using each cell’s volume, which was obtained from the buoyant mass measurement, assuming a cell density of 1.05 g/ml. These data were then filtered according to the following criteria: 1) cells with negative mass accumulation rate (i.e. dying cells) were excluded, 2) only cells within the 30-100 pg size range were included, 3) genes that were expressed/detected in less than 30 L1210 cells or in less than 90 FL5.12 cells were excluded, and 4) cells were excluded when the 30 most abundant transcripts (across all data) were not detected. The remaining dataset consisted of 7994 different mRNAs in 88 L1210 cells and of 8035 different mRNAs in 203 FL5.12 cells. Log-converted mRNA data were then correlated with the log-converted buoyant mass of each cell, and the scaling factor (slope of the correlation) was used to evaluate size scaling. The grouping of genes was done according to GO-terms retrieved from the Mouse Genome Database (MGD), Mouse Genome Informatics, The Jackson Laboratory, Bar Harbor, Maine (http://www.informatics.jax.org, retrieved May, 2022) ^72^. The GO-terms examined were External side of plasma membrane (GO:0009897), and Cell division (GO:0051301).

### SMR data gating and scaling factor analysis

SMR data from each sample was independently gated and analyzed using MATLAB. The first gating step was manually conducted on cell buoyant mass to remove particles too small to be considered cells. The data was then gated for viable cells using a rectangular gating on amine labeling intensity vs buoyant mass (Fig. S2E). Next, a 2D ellipsoid gating (on amine labeling intensity vs buoyant mass) was performed on the live cell population to remove outliers outside of a 95% confidence interval of the population. To compute the ellipsoid gate, we first calculated the covariance matrix of the amine labeling intensity and buoyant mass values. Eigenvectors and eigenvalues of the covariance matrix were then calculated, and the smallest and largest eigenvalues were determined. These values were then used to determine the semi-major and semi-minor axes of the ellipse. The contour of the ellipse was set around the mean value, and axe lengths were determined by parameters for a 95% confidence interval using a chi-square value = 2.4777. Cells within the ellipsoid gate were then used for the scaling factor analysis. For the WGA labeled samples, an additional gating was applied to remove cell aggregates that were specific to this staining approach.

For cell cycle specific analyses of the scaling factor, we used geminin-GFP vs buoyant mass data to differentiate G1 and S&G2 populations with a quadratic gating design (Fig. 2A). For polyploidy analysis, different cell cycle generations were determined by manual polygon gating on geminin-GFP vs buoyant mass, which was done with MATLAB function drawpolygon().

For sensitivity analysis (Fig. S2G), manual gating on amine labeling vs buoyant mass was done with MATLAB function drawpolygon(). For rectangular gating in the sensitivity analysis, the amine labeling intensity of each cell was first normalized by its buoyant mass and the gating was performed on mass-normalized amine label vs buoyant mass. Bootstrap analysis (Fig. S2F) was conducted with resampling with replacement using MATLAB function randi() on the cell indices from the sample population.

The scaling factor was determined by the slope fitting using MATLAB linear regression function regstats(x, y, ‘linear’) on the log_2_ transformed data. The fitted slope and r-square values were then recorded. To compute the 95% confidence interval of the slope, coefCI() function was used.

To analyze how cell cycle transitions influence cell surface amine labeling, we separated cell cycle stages as shown in Fig. 2A, and we identified identically sized cell pairs in different cell cycle stages which were used to compare amine labeling intensity changes between the cell cycle stages. Notably, the separation between G2/M and cytokinetic cells is carried out based on the degradation of geminin-GFP, which separates cells before anaphase onset from those after anaphase onset. The data for L1210 cells is adapted from ^53^. For validations, see ^53^.

### Statistical analyses

All statistical analyses and calculated p-values are detailed in the figures and figure legends. Relevant statistical parameters, including replication information, are also detailed in the figure legends. Results with a p-value below 0.05 were considered significant. For all linear regression analyses, i.e. scaling factor analyses, data linearity was examined visually, but not statistically tested. Statistical calculations were carried out in MATLAB (version 2020b) or OriginPro (version 2023).

## Notes

### Summary of Updates

New experimental data, found in Figures 1, 4, and S4. These data support our original conclusion, but expand the experimental validations to more measured metrics. In addition, the manuscript title, abstract, and discussion have been updated.

