## Supplemental Figures for "Plasma membrane folding enables constant surface area-to-volume ratio in growing mammalian cells"

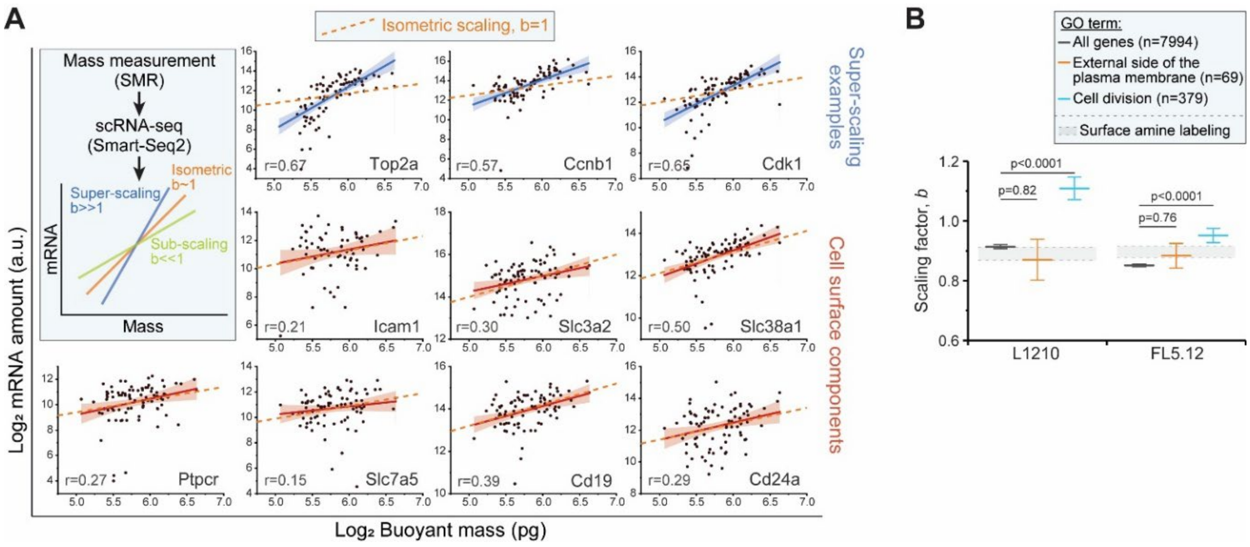


**Figure S1. Coupled buoyant mass and scRNA-seq measurements indicate isometric size-scaling of cell surface associated transcripts. Related to Figure 1.**

**(A)** Scaling of mRNA levels with buoyant mass in single cells. Data obtained from Kimmerling, et al., 2018. SMR-based single-cell mass measurements were combined with scRNA-seq. Scatterplots display examples of L1210 cell mRNA size scaling behaviors for cell cycle associated transcripts (blue, top row) and for cell surface associated transcripts (red, middle and bottom rows). Each dot is a separate cell, line and shaded area represent power law fits and their 95% confidence intervals, and dashed orange lines indicate isometric scaling. Due to the noise in scaling factors of individual mRNAs, cells were grouped based on their GO-term association for future analyses.

**(B)** mRNA scaling factors for transcripts included in the indicated GO-terms in L1210 (n=87 cells) and FL5.12 cells (n=203 cells). Cell surface protein labeling results are shown for reference as a light grey area. p-values were calculated using ANOVA followed by Tukey's posthoc test. Bar and whiskers depict mean±SE.

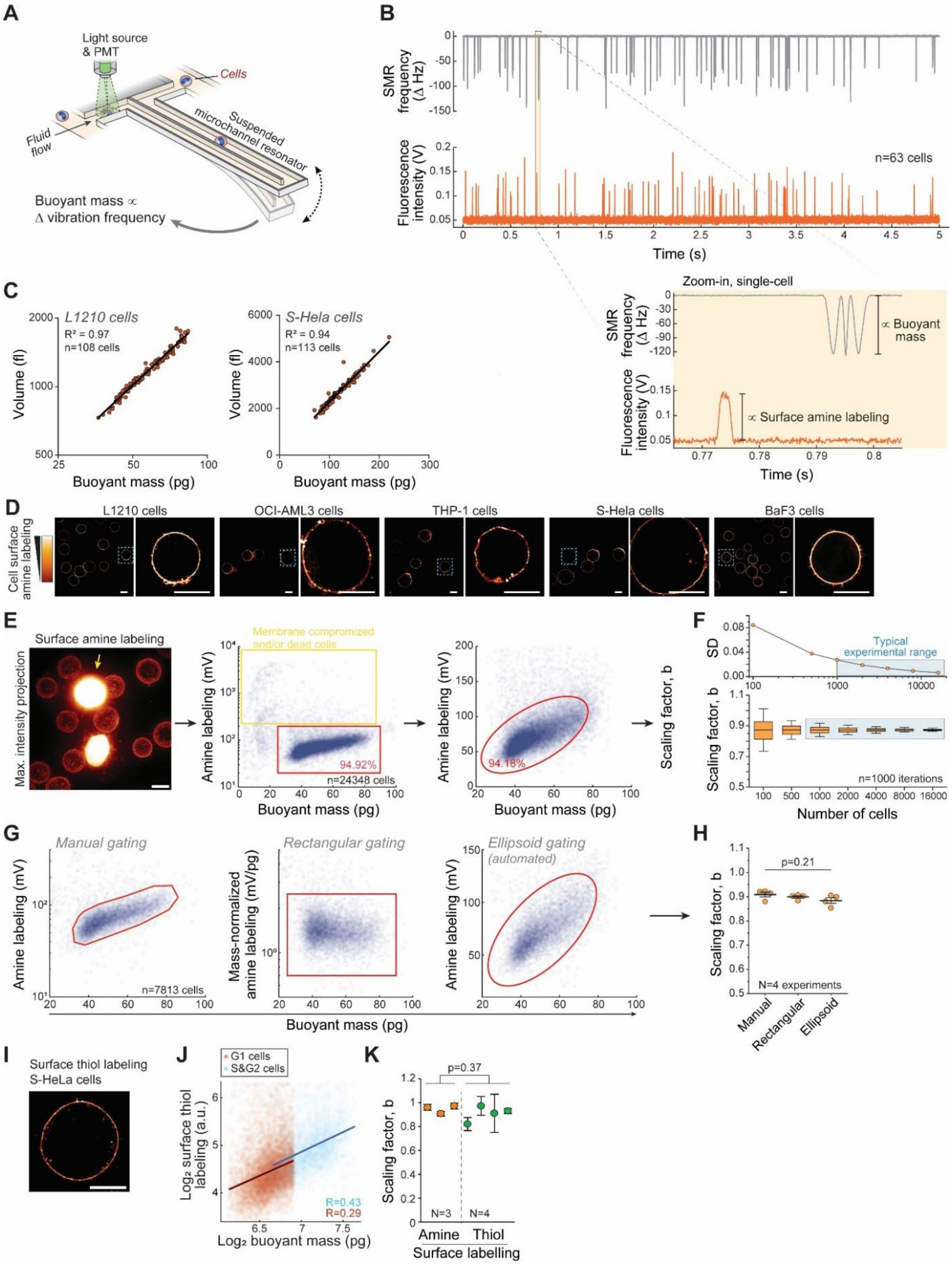


**Figure S2. Measuring the cell size scaling of fluorescent labels using the SMR. Related to Figure 1.**
**(A)** Schematic of the measurement setup. A cell is flown through the SMR, where the cell's buoyant mass is measured based on the change in the SMR's vibration frequency. Immediately next to the SMR cantilever, the cell

passes by a fluorescence measurement area, where fluorescence markers are detected using photomultiplier tubes (PMTs).

**(B)** Representative SMR and PMT raw data of L1210 cell buoyant mass (*top*) and fluorescent cell surface protein labeling (*bottom*) measurements. Inset (*bottom right*) displays a zoom-in to the raw data of a single cell.

**(C)** Representative experiments of L1210 (*left*) and S-HeLa (*right*) cell volumes as a function of buoyant mass. Each dot is a single cell, and the solid line indicates a linear correlation. These results are consistent with our previous analyses showing high correlations between cell volume and buoyant mass in 5 out of the 6 cell lines studied in this manuscript (typical $R^2$ = 0.94, Wu et al., 2024).

**(D)** Representative single z-slice microscopy images of cell surface amine labeling in indicated cell lines. L1210 and BaF3 cell zoom-in example is the same as in Fig. 1. Scale bars denote 10 µm.

**(E)** *Left,* a maximum intensity image of surface amine labelled L1210 cells, where excessively bright cells (yellow arrow) have compromised membrane integrity (i.e. dead cells that are labeled internally). Scale bar denotes 10 µm. *Middle,* representative raw data of L1210 cell surface amine labeling as a function of buoyant mass. Each opaque dot depicts a single cell. The region of viable cells (red gate). *Right,* the viable cells were further gated using an automated ellipsoid gating for the scaling factor analysis.

**(F)** Bootstrapping analysis of scaling factors. *Top*, the standard deviation of the scaling factors as a function of cells analyzed.

**(G)** Examples of three different data gating approaches.

**(H)** Cell surface protein scaling factors derived using the gating approaches indicated in panel (G). Each dot represents a separate experiment, line and whiskers indicate mean±SD. p-value was obtained using ANOVA.

**(I)** A single z-slice image of S-Hela cell labelled for cell surface proteins using a thiol reactive labeling approach.

**(J)** The scaling between cell mass and surface thiol labeling in S-HeLa cells. Each opaque point represents a single cell in G1 (red) or S&G2 (blue) cell cycle stage (n>1000 cells).

**(K)** Comparison of scaling factors obtained with cell surface amine and thiol labeling in S-HeLa cells. Each dot represents a separate experiment. p-value was obtained using Welch's t-test.

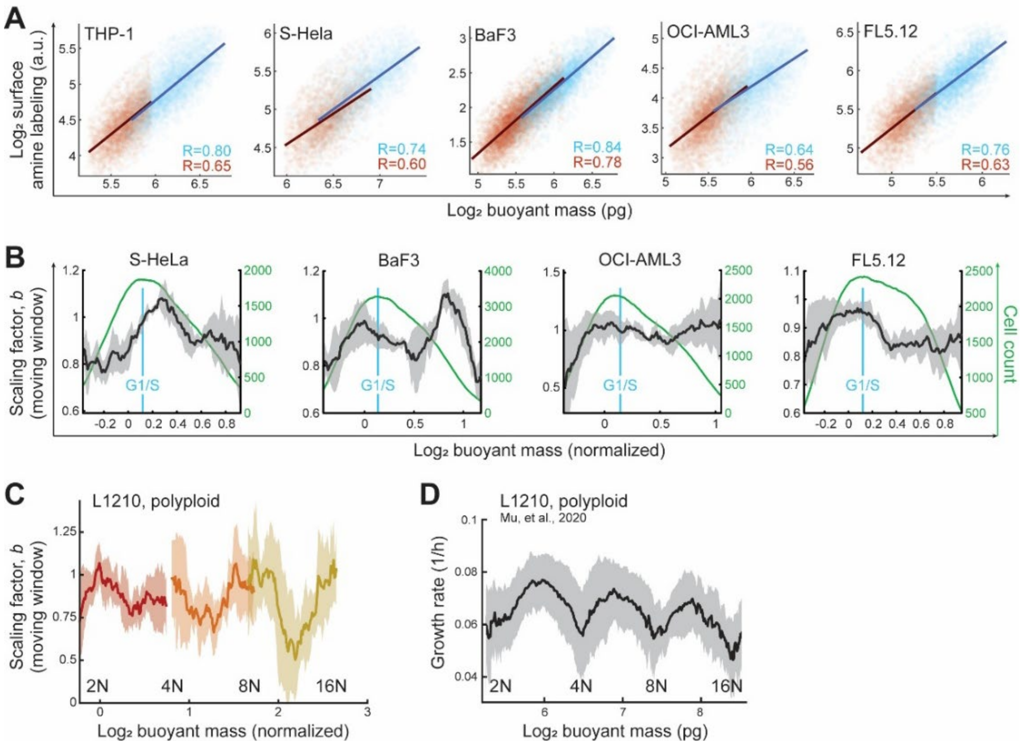


**Figure S3. Cell cycle-dependency of cell surface protein content. Related to Figure 2.**

**(A)** Representative scatter plots displaying the scaling between cell surface protein labeling and cell mass in G1 (red) and S&G2 (blue) stages in indicated cell lines. Each opaque point represents a single cell, lines represent power law fits to each cell cycle stage (n>500 cells per cell cycle stage).

**(B)** Moving window analysis of the scaling factor, $b$, as a function of cell mass in indicated cell lines. Dark line and shaded areas represent the mean±SD of the scaling factor (N=4 independent experiments), green line represents cell size distribution.

**(C)** Moving window analysis of the scaling factor, $b$, as a function of cell mass in L1210 cells of indicated ploidy. Lines and shaded areas represent the mean±SD of the scaling factor, which was analyzed separately for each ploidy (N=9 independent experiments).

**(D)** Mass-normalized L1210 cell growth rates as L1210 cells grow into polyploidy. Data obtained from Mu, et al., 2020. Data represents mean±SD of 76 independent single-cell growth traces.

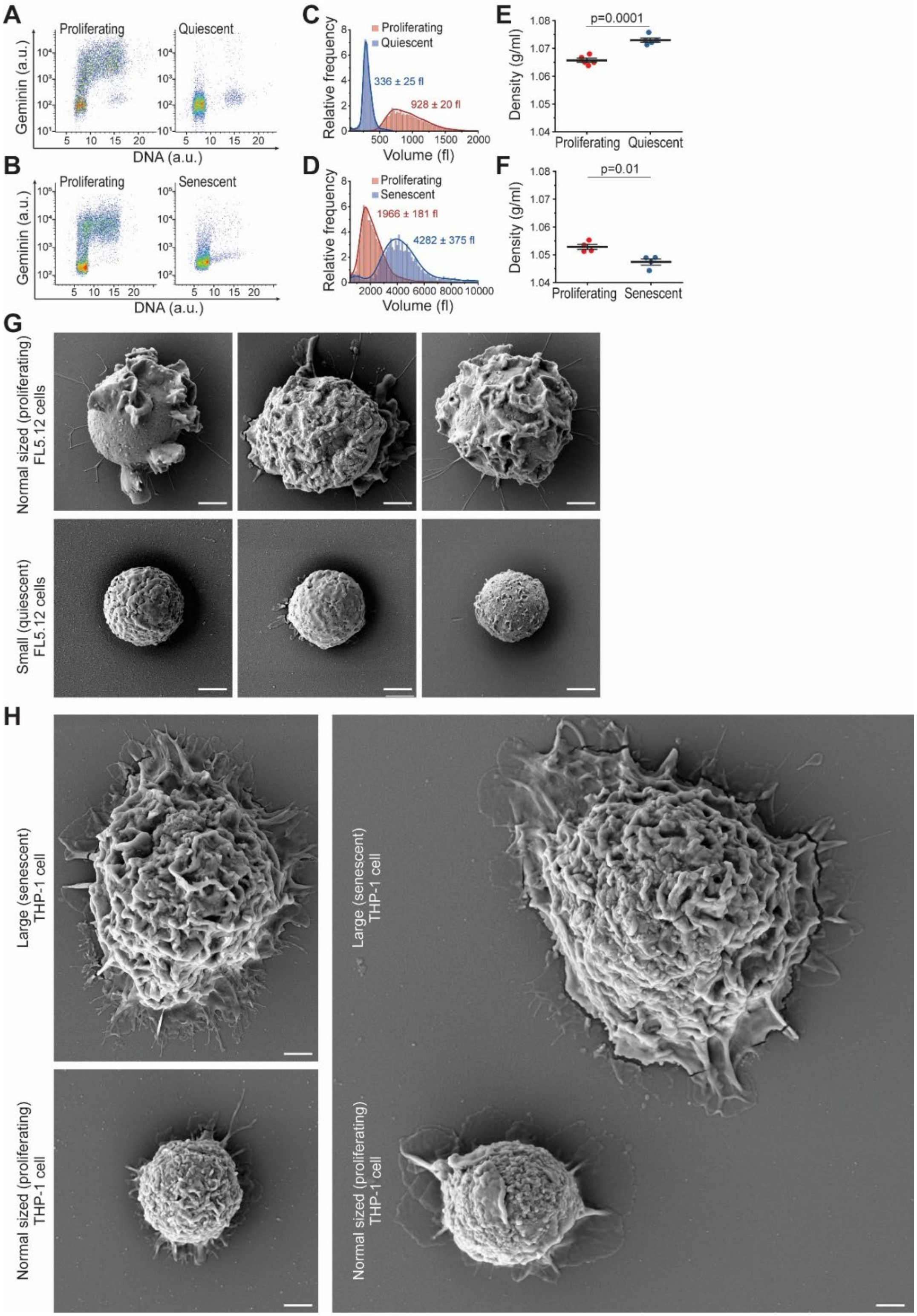


**Figure S4. Cell cycle, cell size, and plasma membrane characterization of quiescent and senescent cell models. Related to Figure 4.**

**(A-B)** Flow cytometry analysis of Geminin-GFP as a function of DNA content in proliferating and non-proliferating FL5.12 cells (A) and THP-1 cells (B). N=4 independent experiments.

**(C-D)** Representative cell volume histograms of proliferating and non-proliferating FL5.12 cells (C) and THP-1 cells (D). Colored texts represent cell volume mean±SD (N=4-5 independent experiments).
**(E-F)** Quantifications of average buoyant densities in populations of proliferating and non-proliferating FL5.12 cells (E) and THP-1 cells (F). Each dot represents a separate experiment (N=4-5), bar and whiskers represent mean±SD, and p-values were obtained using Welch's t-test.
**(G)** Representative SEM images of normal-sized (proliferating) and small (quiescent) FL5.12 cells (N=3 independent experiments, n=70 cells).
**(H)** Representative SEM images of normal-sized (proliferating) and large (senescent) THP-1 cells (N=2 independent experiments, n=25 cells).
All scale bars denote 2 μm.

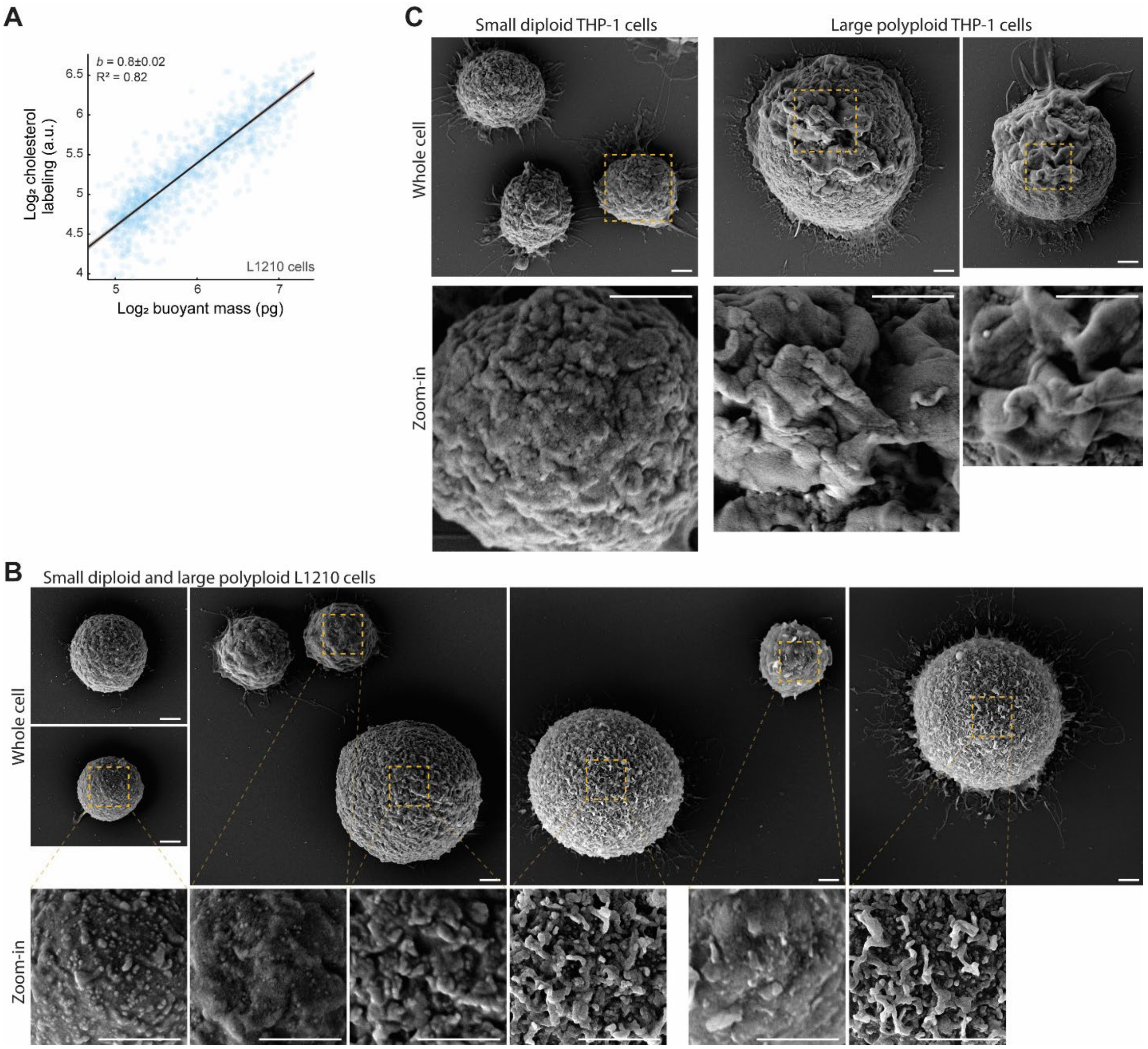


**Figure S5. Plasma membrane folding increases upon excessive cell size increases. Related to Figure 5.**
**(A)** A representative scatter plot displaying the scaling between cell mass and Filipin III cholesterol labeling in L1210 cells. Each opaque point represents a single cell (n=1312 cells). Line and shaded areas represent a power law fit and its 95% confidence intervals.
**(B)** Additional examples of SEM images of small diploid and large polyploid L1210 cells, as in Fig. 5H.
**(C)** Representative SEM images of small diploid and large polyploid THP-1 cells (N=2 independent experiments, n=31 cells).
All scale bars denote 2 μm.
